# Hemimetabolous genomes reveal molecular basis of termite eusociality

**DOI:** 10.1101/181909

**Authors:** Mark C Harrison, Evelien Jongepier, Hugh M. Robertson, Nicolas Arning, Tristan Bitard-Feildel, Hsu Chao, Christopher P. Childers, Huyen Dinh, Harshavardhan Doddapaneni, Shannon Dugan, Johannes Gowin, Carolin Greiner, Yi Han, Haofu Hu, Daniel S.T. Hughes, Ann-Kathrin Huylmans, Carsten Kemena, Lukas P.M. Kremer, Sandra L. Lee, Alberto Lopez-Ezquerra, Ludovic Mallet, Jose M. Monroy-Kuhn, Annabell Moser, Shwetha C. Murali, Donna M. Muzny, Saria Otani, Maria-Dolors Piulachs, Monica Poelchau, Jiaxin Qu, Florentine Schaub, Ayako Wada-Katsumata, Kim C. Worley, Qiaolin Xie, Guillem Ylla, Michael Poulsen, Richard A. Gibbs, Coby Schal, Stephen Richards, Xavier Belles, Judith Korb, Erich Bornberg-Bauer

**Author notes:** Corresponding authors. (XB); (JK); (EBB). These authors contributed equally to this work (Mark C Harrison, Evelien Jongepier, Hugh M. Robertson).

## Abstract

Around 150 million years ago, eusocial termites evolved from within the cockroaches, 50 million years before eusocial Hymenoptera, such as bees and ants, appeared. Here, we report the first, 2GB genome of a cockroach, *Blattella germanica*, and the 1.3GB genome of the drywood termite, *Cryptotermes secundus*. We show evolutionary signatures of termite eusociality by comparing the genomes and transcriptomes of three termites and the cockroach against the background of 16 other eusocial and non-eusocial insects. Dramatic adaptive changes in genes underlying the production and perception of pheromones confirm the importance of chemical communication in the termites. These are accompanied by major changes in gene regulation and the molecular evolution of caste determination. Many of these results parallel molecular mechanisms of eusocial evolution in Hymenoptera. However, the specific solutions are remarkably different, thus revealing a striking case of convergence in one of the major evolutionary transitions in biological complexity.

---

Eusociality, the reproductive division of labour with overlapping generations and cooperative brood care, is one of the major evolutionary transitions in biology.^1^ Although rare, eusociality has been observed in a diverse range of organisms, including shrimps, mole-rats and several insect lineages.^2, 3, 4^ A particularly striking case of convergent evolution occurred within the holometabolous Hymenoptera and in the hemimetabolous termites (Isoptera), which are separated by over 350 my of evolution.^5^ Termites evolved within the cockroaches around 150 mya, towards the end of the Jurassic,^6,7^ about 50 my before the first bees and ants appeared.^5^ Therefore, identifying the molecular mechanisms common to both origins of eusociality is crucial to understanding the fundamental signatures of these rare evolutionary transitions. While the availability of genomes from many eusocial and non-eusocial hymenopteran species^8^ has allowed extensive research into the origins of eusociality within ants and bees,^9,10,11^ a paucity of genomic data from cockroaches and termites has precluded large-scale investigations into the evolution of eusociality in this hemimetabolous clade.

The conditions under which eusociality arose differ greatly between the two groups. Termites and cockroaches are hemimetabolous and so show a direct development, while holometabolous hymenopterans complete the adult body plan during metamorphosis. In termites, workers are immatures and only reproductive castes are adults,^12^ while in Hymenoptera, adult workers and queens represent the primary division of labour. Moreover, termites are diploid and their colonies consist of both male and female workers, and usually a queen and king dominate reproduction. This is in contrast to the haplodiploid system found in Hymenoptera, in which all workers and dominant reproductives are female. It is therefore intriguing that strong similarities have evolved convergently within the termites and the hymenopterans, such as differentiated castes and a nest life with reproductive division of labour. The termites can be subdivided into wood-dwelling and foraging termites. The former belong to the lower termites and produce simple, small colonies with totipotent workers that can become reproductives. Foraging termites (some lower and all higher termites) form large, complex societies, in which worker castes can be irreversible.^12^ For this reason, higher, but not lower, termites can be classed as superorganismal.^13^ Similarly, within Hymenoptera, varying levels of eusociality exist.

Here we provide insights into the molecular signatures of eusociality within the termites. We analysed the genomes of two lower and one higher termite species and compared them to the first cockroach genome, as a closely-related non-eusocial outgroup. Furthermore, differences in expression between nymphs and adults of the cockroach were compared to differences in expression between workers and reproductives of the three termites, in order to gain insights into how expression patterns changed along with the evolution of castes. Using fifteen additional insect genomes to infer background gene family turnover rates, we analysed the evolution of gene families along the transition from non-social cockroaches to eusociality in the termites. In this study we concentrated particularly on two hallmarks of insect eusociality, as previously described for Hymenoptera, with the expectation that similar patterns occurred along with the emergence of termites. These are the evolution of a sophisticated chemical communication, which is essential for the functioning of a eusocial insect colony^3, 14, 15^ and major changes in gene regulation along with the evolution of castes.^9, 10^ Additionally, we tested if transposable elements spurred the evolution of gene families that were essential for the transition to eusociality.

## Evolution of genomes, proteomes and transcriptomes

We sequenced and assembled the genome of the German cockroach, *Blattella germanica* (Ectobiidae), and of the lower, drywood termite, *Cryptotermes secundus* (Kalotermitidae; for assembly statistics see Supplementary Table 1). The cockroach genome (2.0 Gb) is considerably larger than all three termite genomes. The genome size of *C. secundus* (1.30 Gb) is comparable to the higher, fungus-growing termite, *Macrotermes natalensis*, (1.31 Gb, Termitidae)^16^ but more than twice as large as the lower, dampwood termite, *Zootermopsis nevadensis* (562 Mb, Termopsidae).^17^ The smaller genomes of termites compared to the cockroach are in line with previous size estimations based on C-values.^18^ The proteome of *B. germanica* (29,216 proteins) is also much larger than in the termites, where we find the proteome size in *C. secundus* (18,162) to be similar to the other two termites (*M. natalensis*: 16,140; *Z. nevadensis*: 15,459; Fig. 1). In fact, the *B. germanica* proteome was the largest among all 21 arthropod species analysed here (Fig. 1). Strong evidential support for over 80% of these proteins in *B. germanica* (see Supplementary Material) and large expansions in many manually annotated gene families offer high confidence in the accuracy of this proteome size.

**Figure 1.**
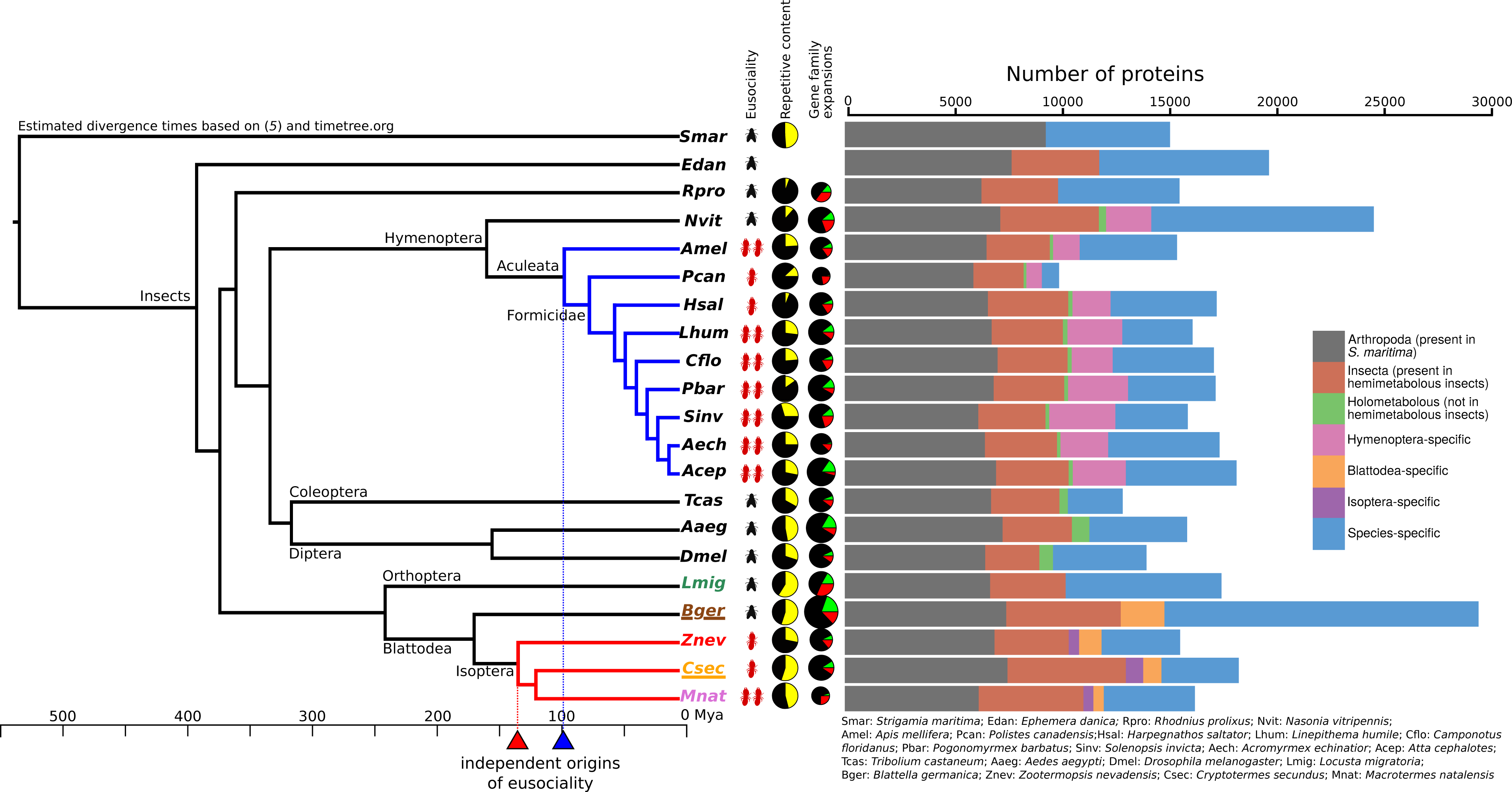
Phylogenetic, genomic and proteomic comparisons of 20 insect species. From left to right: Phylogenetic tree of 20 insect species with *Strigamia maritima* (centipede) as outgroup; level of eusociality (one red insect: simple eusociality; two red insects: advanced eusociality; black fly: non-eusocial); fractions of repetitive content (yellow) within genomes of selected species (for sources see supplementary material); proportions of species-specific gene family expansions (green), contractions (red) and stable gene families (black), size of pies represents relative size of gene family change (based on total numbers). Bar chart showing protein orthology across taxonomic groups within each genome.

We also compared gene expression between the four species. When comparing worker expression with queen expression in the termites and nymph expression (5th and 6th instars) with adult female expression in *B. germanica*, we found shifts in specificity of expression for termites compared to the cockroach in several gene families (Fig. 2). It has been previously reported for the primitively eusocial paper wasp, *Polistes canadensis*, that the majority of caste-biased genes, especially those upregulated in workers, are novel genes.^19^ The authors suggested that this may be a feature of early eusociality. We did not find the same pattern for the termites. Species-specific genes (those without ortholog) were not enriched for differentially expressed genes in any of the termites, with slight peaks among Blattodea- and Isoptera-specific genes (Supplementary Figure 1).

**Figure 2.**
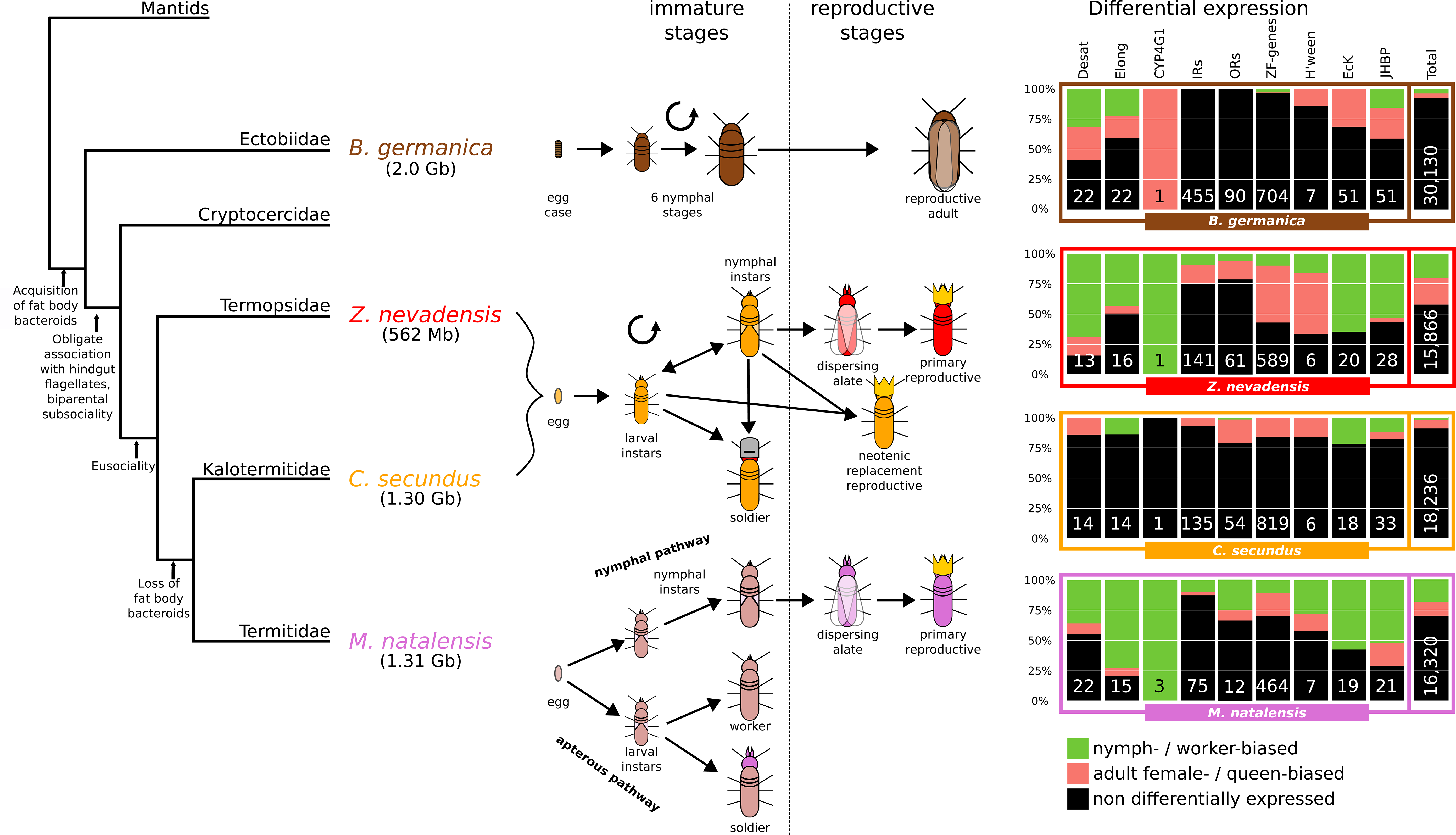
Comparison of developmental pathways between *B. germanica*, the lower termites, *Z. nevadensis* and *C. secundus*, and the higher termite, *M. natalensis.* Shown from left to right are: a simple phylogeny^97^ describing important novelties along the evolutionary trajectory to termites (numbers in brackets are genome sizes); life cycles; differential expression (log_2_FoldChange > 1 & p < 0.05) between workers and queens (between nymphs and adult females in B.*germanica*) of selected gene families (Desat = desaturases, Elong = elongases, H’ween = Halloween genes) and total numbers within all genes; numbers denote total numbers of genes in each gene family.

## Gene family expansions assisted by TEs in termites

The transitions to eusociality in ants^10^ and bees^9^ have been linked to major changes in gene family sizes. Similarly, we detected significant gene family changes on the branch leading to the termites (7 expansions and 10 contractions; Supplementary Figure 2, Supplementary Table 2). The numbers of species-specific, significant expansions and contractions of gene families varied within termites (*Z. nevadensis*: 15/5; *C. secundus*: 27/3; *M. natalensis*: 24/20; Supplementary Figure 2 & Supplementary Tables 3-5). Interestingly, in *B. germanica* we measured 93 significant gene family expansions but no contractions (Supplementary Table 6), which contributed to the large proteome.

The termite and cockroach genomes contain a higher level of repetitive DNA compared to the hymenopterans we analysed (Fig. 1). *C. secundus* and *B. germanica* genomes both contain 55% repetitive content (Supplementary Table S7), which is higher than in both *Z. nevadensis* (28%) and the higher termite, *M. natalensis* (46%; Fig. 1).^20^ As also found in *Z. nevadensis* and *M. natalensis*,^20^ LINEs and especially the subfamily BovB were the most abundant transposable elements (TEs) in the *B. germanica* and *C. secundus* genomes, indicating that a proliferation of LINEs may have occurred in the ancestors of Blattodea (cockroaches and termites).

We hypothesised that these high levels of TEs may be driving the high turnover in gene family sizes within the termites and *B. germanica*.^21^ Expanded gene families indeed had more repetitive content within 10 kb flanking regions in all three termites (p < 1.3x10*^−8^*; Wald *t*-test; Supplementary Tables 8-9), in particular in the higher termite *M. natalensis*. In contrast, gene family expansions were not correlated with TE content in flanking regions for *B. germanica*. These results suggest a major expansion of LINEs at the root of the Blattodea clade contributed to the evolution of gene families within termites, likely via unequal crossing-over;^21^ however, the expansions in *B. germanica* were not facilitated by TEs. It can therefore be speculated that the large expansion of LINEs within Blattodea allowed the evolution of gene families which ultimately facilitated the transition to eusociality.

## Massive expansion and positive selection of Ionotropic Receptors

Insects perceive chemical cues from toxins, pathogens, food and pheromones with three major families of chemoreceptors, the Odorant (ORs), Gustatory (GRs) and Ionotropic (IRs) Receptors.^22^ Especially ORs have been linked to colony communication in eusocial Hymenoptera, where they abound.^14, 15, 23^ Interestingly, as previously detected for *Z. nevadensis*,^17^ the OR repertoire is substantially smaller in *B. germanica* and all three termites compared to hymenopterans. IRs, on the other hand, which are less frequent in hymenopterans, are strongly expanded in the cockroach and termite genomes (Fig. 3 & Supplementary Figure 3). Intronless IRs, which are known to be particularly divergent,^24^ show the greatest cockroach and Blattodea-specific expansions (Fig. 3a, Blattodea-, Cockroachand Group D-IRs). By far the most IRs among all investigated species were found in *B. germanica* (455 complete gene models), underlining that the capacity for detecting many different kinds of chemosensory cues is crucial for this generalist that thrives in challenging, human environments. In line with a specialisation in diet and habitat, the total number of IRs is lower within the termites (*Z. nevadensis*: 141; *C. secundus*: 135; *M. natalensis*: 75). Nevertheless, IRs are more numerous in termites than in all other analysed species (except *Nasonia vitripennis*: 111). This is strikingly similar to the pattern for ORs in Hymenoptera, which are also highly numerous in non-eusocial outgroups as well as in eusocial species.^14, 23, 25^.

**Figure 3.**
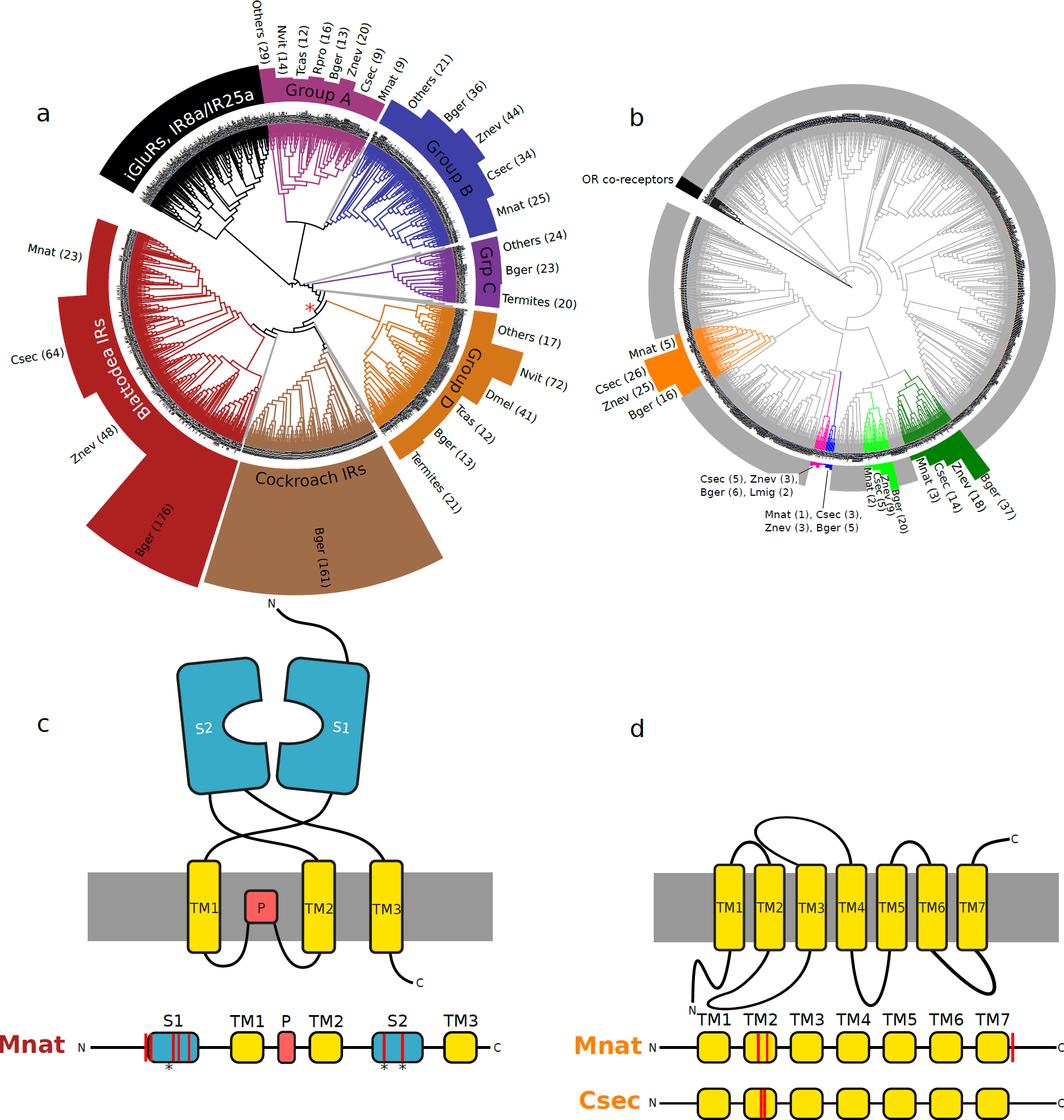
Expansions, contractions and positive selection within IRs and ORs in termites. **a.**IR and **b.** OR gene trees of 13 insect species. In each tree only well supported clades (support values *>* 85) that include *B. germanica* or termite genes are highlighted within the gene trees. Lengths of coloured bars represent number of genes per species within each of these clades. Red asterisk in a**b.** denotes putative root of intronless IRs. **c.** The upper cartoon depicts the 2D structure of an IR, containing ligand binding lobes (S1 & S2), transmembrane regions (TM1-3) and the pore domain (P). Below, the sequence of the domains along the peptide is represented, showing that the sites, which are under significant positive selection (red bars; codeml site models 7 & 8) within Blattodea-IRs for *M. natalensis* (p < 1.7x10*^−10^*) are all situated within the ligand binding lobes and on or around the putative ligand binding sites (asterisks).^86^ **d.** The same representation for ORs, which include 8 transmembrane regions. Positive selection was found for *M. natalensis* (p = 1.1x10*^−11^*) and *C. secundus* (p = 5.6x10*^−16^*) of the orange clade, each at two codon positions within the second transmembrane region and at a third position within the C-terminal extra-cellular region for *M. natalensis*.

We scanned each IR group for signs of species-specific positive selection. Within the Blattodea-specific intronless IRs, we found several codon positions under significant positive selection for the higher termite, *M. natalensis* (codeml site models 7 & 8; p < 1.7x10*^−10^*). The positively evolving codons are situated within the two ligand-binding lobes of the receptors (Fig. 3c), showing that a diversification of ligand specificity has occurred along with the transition to higher eusociality and a change from wood-feeding to fungus-farming in *M. natalensis*. Only two IRs were differentially expressed between nymphs and adult females in *B. germanica*. Underlining a change in expression along with the evolution of castes, we found 35 IRs to be differentially expressed between workers and queens in *Z. nevadensis*, 11 in *C. secundus* and 10 in *M. natalensis* (Fig. 2, Supplementary Table 10). The possible role of IRs in pheromonal communication has been highlighted both in the cockroach *Periplaneta americana*^26^ and in *Drosophila melanogaster*,^27^ where several IRs show sex-biased expression.

One group of ORs (orange clade in Fig. 3b) is evolving under significant positive selection at codon positions within the second transmembrane domain in *M. natalensis* (codeml site model; p = 1.1x10*^−11^*) and *C. secundus* (p = 5.6x10*^−16^*; Fig. 3d). Such a variation in the transmembrane domain can be related to ligand binding specificity, as has been shown for a polymorphism in the third transmembrane domain for an OR in *D. melanogaster*,^28, 29^ adding further support for an adaptive evolution of chemoreceptors, in line with the greater need for a sophisticated colony communication in the termites. Similar to IRs, a higher proportion of ORs were differentially expressed between workers and queens in the three termites than between nymphs and adults in the cockroach (Fig. 2; Supplementary Table 11), highlighting a change in expression and function along with the transition to eusociality. The evolution of chemoreceptors along with the emergence of the termites can also be related to adaptation processes and changes in diet compared to the cockroach. Experimental verification will help pinpoint which receptors are particularly important for communication.

## CHC producing enzymes have evolved caste-specificity

Despite their different ancestry, both termites and eusocial hymenopterans are characterised by the production of caste-specific cuticular hydrocarbons (CHCs),^30, 31, 32^ which are often crucial for regulating reproductive division of labour and chemical communication. Accordingly, we find changes in the termites in three groups of proteins involved in the synthesis of CHCs: desaturases (introduction of double bonds^33^), elongases (extension of C-chain length^34^) and CYP4G1 (last step of CHC biosynthesis^35^).

Desaturases are thought to be important for division of labour and social communication in ants.^36^As previously described for ants,^36^ Desat B genes are the most abundant desaturase family in the termites and the cockroach (Supplementary Table 12), especially in *M. natalensis* where we found ten gene copies (significant expansion; p = 0.0003; Supplementary Table 5; Supplementary Figure 4). As in ants, especially the First Desaturases (Desat A Desat E) vary greatly in their expression between castes and species in the three termites (Fig. 2; Supplementary Table 13).^36^ In contrast to ants, where these genes are under strong purifying selection,^36^ we found significant positive selection within the Desat B genes for the highly eusocial termite, *M. natalensis*, (codeml site models 7 & 8; p = 1.1x10*^−16^*), indicating a diversification in function, possibly related to their greater diversification of worker castes (major and minor workers, major and minor soldiers). Although desaturases are often discussed in the context of CHC production and chemical communication, their biochemical roles are quite diverse,^36^ and the positive selection we observe for *M. natalensis* may, at least in part, be related to their rather different ecology of foraging and fungus farming rather than nest mate recognition. Future experimental verification of the function of these genes will help better understand these observed genomic and transcriptomic patterns.

Underlining an increased importance of CHC communication in termites, the expression patterns of elongases (extension of C-chain length) differ considerably in the termites compared to the cockroach (Fig. 2; Supplementary Table 14). In contrast to *B.germanica*, in which elongases are both nymph(5 genes) and adult-biased (4 genes), only one or two elongase genes in each termite are queen-biased in their expression, while many are worker-biased. As with the desaturases, a group of *M. natalensis* elongases also reveal significant signals of positive selection (codeml branch-site test; p = 4x10*^−4^*), further indicating a greater diversification of CHC production in this higher termite.

The last step of CHC biosynthesis, the production of hydrocarbons from long-chain fatty aldehydes, is catalyzed by a P450 gene, CYP4G1, in *D. melanogaster*.^35^ We found one copy of CYP4G1 in *B. germanica*, *Z. nevadensis* and *C. secundus*, but three copies in *M. natalensis*, reinforcing the greater importance of CHC synthesis in this higher termite. Corroborating the known importance of maternal CHCs in *B. germanica*,^37^ CYP4G1 is over-expressed in female adults compared to nymphs (Fig. 2; Supplementary Table 15). In each of the termites, however, CYP4G1 is more highly expressed in workers (or kings in *C. secundus*) compared to queens (Fig. 2; Supplementary Table 15), adding support that, compared to cockroach nymphs, a change in the dynamics and turnover of CHCs in termite workers has taken place.

## Changes in gene regulation in termites

The development of distinct castes underlying division of labour is achieved via differential gene expression. Major changes in gene regulation have been reported as being central to the transition to eusociality in bees^9^ and ants.^10^ Accordingly, we found major changes in putative DNA methylation patterns (levels per 1-to-1 ortholog) among the termites compared to four other hemimetabolous insect species (Fig. 4a). This is revealed by CpG depletion patterns (CpG*_o/e_*), a reliable predictor of DNA methylation,^38, 39^ correlating more strongly between the termites than among any of the other analysed hemimetabolous insects (Fig. 4). In other words, within orthologous genes, predicted DNA methylation levels differ greatly between termites and other hemimetabolous species but remain conserved among termite species.

**Figure 4.**
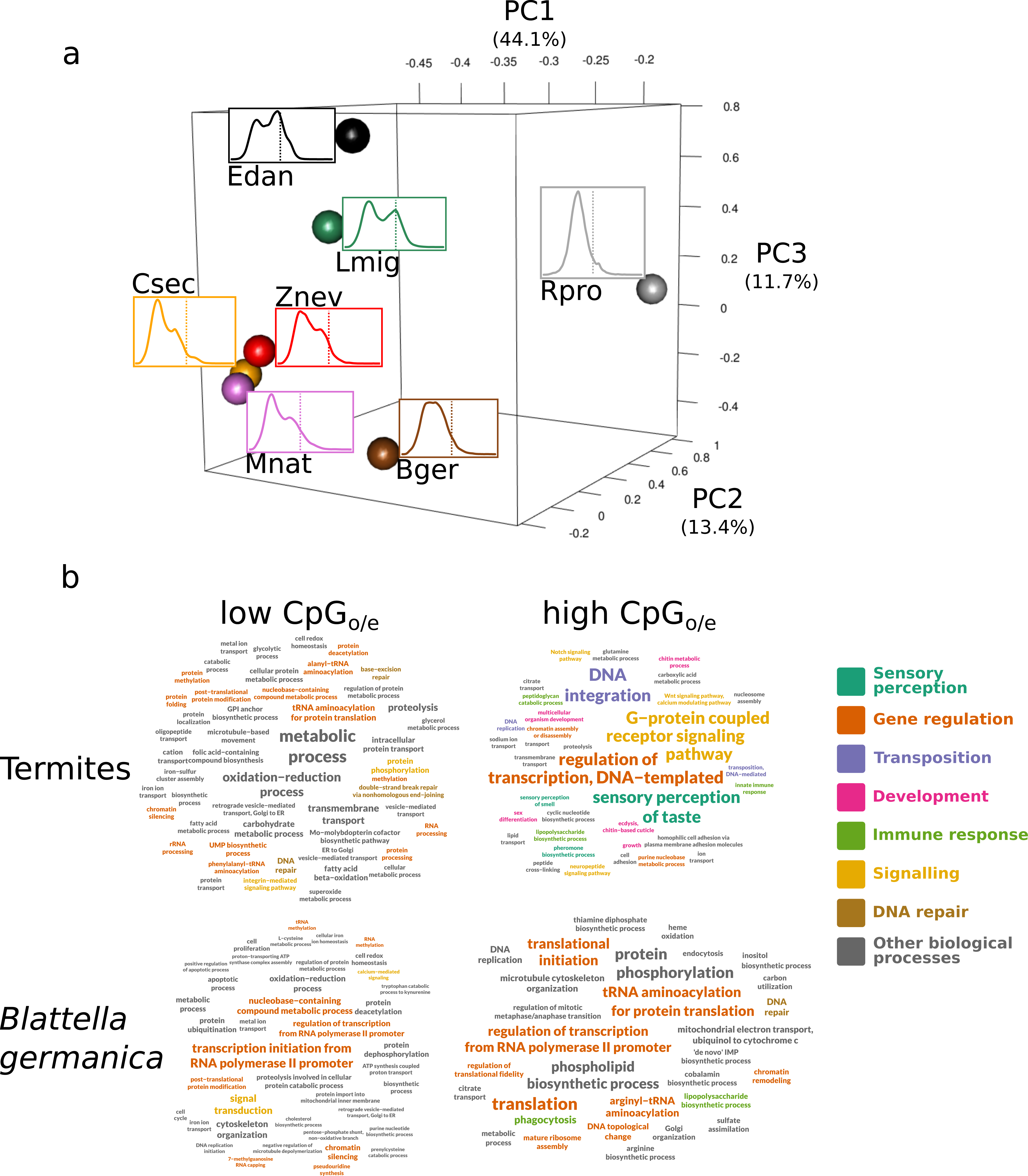
CpG***_o/e_***of seven hemimetabolous insects. **a.** PCA of predicted DNA methylation patterns among 2664 1-to-1 orthologs, estimated via CpG*_o/e_*. Spheres represent positions of species within 3D PCA, with the distance between spheres representing the similarity of CpG*_o/e_* between species at each ortholog; curves are distribution of CpG*_o/e_* with dotted line showing CpG*_o/e_* = 1. **b.** Tag clouds of enriched (p < 0.05) GO terms (biological processes) among lower (left) and higher quartile (right) of CpG*_o/e_* within termites (top) and *B. germanica* (bottom). For termites, genes were merged from all three species for analysing GO term enrichment. High CpG*_o/e_* indicates low level of DNA methylation and vice versa.

Predicted levels of DNA methylation correlated negatively with caste-specificity of expression for each of the termites. This is confirmed by a positive correlation between CpG*_o/e_* (negative association with level of DNA methylation) and absolute log_2_-fold change of expression between queens and workers (Pearson’s r = 0.32 to 0.36; p < 2.2x10*^−16^*). The caste-specific expression of putatively unmethylated genes in termites is reflected in the enrichment of GO terms related to sensory perception, regulation of transcription, signalling and development, whereas methylated genes are mainly related to general metabolic processes (Fig. 4b, Supplementary Table 16). These results show strong parallels to findings for eusocial Hymenoptera.^40, 41, 42, 43^ This is in stark contrast to the non-eusocial cockroach, *B. germanica*, where there was only a very weak relationship between CpG*_o/e_*and differential expression between nymphs and adult females (r = 0.14), nor were any large differences apparent in enriched GO terms between putatively methylated and non-methylated genes (Fig. 4b).

Our results argue in favour of a diminished role of DNA methylation in caste-specific expression within eusocial insects, as recently shown.^38, 44^ In fact, DNA methylation appears to be important for the regulation of house-keeping genes because predicted methylated genes are related to general biological processes (further supported by lower CpG*_o/e_* within 1-to-1 orthologs than in non-conserved genes),^45^ while caste-specific genes are ’released’ from this type of gene regulation. However, a recent study linked caste-specific DNA methylation to alternative splicing in *Z. nevadensis*^46^.

Major biological transitions are often accompanied by expansions of transcription factor (TF) families, such as genes containing zinc-finger (ZF) domains.^47^ We also observed large differences in ZF families within the termites compared to *B. germanica*. Many ZF families were reduced or absent in termites, while different, unrelated ZF gene families were significantly expanded (Supplementary Tables 2-6). Queen-biased genes were significantly over-represented among ZF genes for each of the termites (p < 2 x 10*^−10^*;*χ*^2^ test; Supplementary Table 17), indicating an important role of ZF genes in the regulation of genes related to caste-specific tasks and colony organisation in the termites. This is in contrast to the significant under-representation of differentially expressed ZF genes within *B. germanica* (p = 4.8 x 10*^−5^*; *χ*^2^-test). Interestingly, two other important TF families (bHLH and bZIP),^47^ which were not expanded in the termites, showed no caste-specific expression pattern (p > 0.05), except bZIP genes, in which queen-biased genes were marginally over-represented for *M. natalensis* (p = 0.049). These major upheavals in ZF gene families and their caste-specific expression show that major changes in TFs accompanied the evolution of termites, strikingly similar to the evolution of ants.^10^

## Evolution of genes related to molting and metamorphosis

Hemimetabolous eusociality is characterised by differentiated castes, which represent different developmental stages. This is in contrast to eusocial Hymenoptera, in which workers and reproductives are adults. While cockroaches develop directly through several nymphal stages before becoming reproductive adults, termite development is more phenotypically plastic, and workers are essentially immatures (Fig. 2). In wood-dwelling termites, such as *C. secundus* and *Z. nevadensis*, worker castes are non-reproductive immatures that are totipotent to develop into other castes, while in the higher termite, *M. natalensis*, workers can be irreversibly defined instars. It is therefore clear that a major change during the evolution of termites occurred within developmental pathways. Accordingly, we found changes in expression and gene family size of several genes related both to molting and metamorphosis.

In the synthesis of the molting hormone, 20-hydroxyecdysone, the six Halloween genes (5 Cytochrome P450s and a Rieske-domain oxygenase) play a key role.^48, 49^ Only one Halloween gene, Shade (Shd; CYP314A1), which mediates the final step of 20-hydroxyecdysone synthesis, is differentially expressed between the final nymphal stages and adults females in *B. germanica* (Fig. 2; Supplementary Table 18), consistent with its role in the nymphal or imaginal molt. In the three termites, the Halloween genes show varying caste-specific expression (Fig. 2; Supplementary Table 18), showing that ecdysone plays a significant role in the regulation of caste differences. Ecdysteroid kinase genes (EcK), which convert the insect molting hormone into its inactive state, ecdysone 22-phosphate, for storage,^50^ are only overexpressed in female adults compared to nymphs in *B. germanica* (16/51 genes, Fig.2, Supplementary Table 19). In termites, however, where the gene copy number is reduced (18 to 20 per species), these important molting genes appear to have evolved worker-specific functions (Fig. 2; Supplementary Table 19).

Whereas 20-hydroxyecdysone promotes molting, juvenile hormone (JH) represses imaginal development in pre-adult instars.^51^ JH is important in caste differentiation in eusocial insects, including termites.^12, 52^ Hemolymph juvenile hormone binding proteins (JHBP), which transport JH to its target tissues,^53^ are reduced within the termites (21 to 33 genes) but significantly expanded in *B. germanica* (51 copies; p = 0018; Supplementary Table 6). Thirteen of the JHBP genes are over-expressed in adult females and only 8 in nymphs in *B. germanica* (Fig. 2, Supplementary Table 20). In both *Z. nevadensis* and *M. natalensis*, on the other hand, JHBPs are significantly more worker-biased (p < 0.01, *χ*^2^ test; Supplementary Table 20; Fig. 2). In *C. secundus*, expression is more varied, with 4 worker-biased, 7 king-biased and 2 queen-biased genes (Fig. 2; Supplementary Table 20).

These changes in copy number and caste-specific expression of genes involved in molting and metamorphosis within termites compared to the German cockroach demonstrate that changes occurred in the control of the developmental pathway along with the evolution of castes. However, this interpretation needs to be experimentally verified.

## Conclusions

These results, considered alongside many studies on eusociality in Hymenoptera,^9, 10, 14, 36^ provide evidence that major changes in gene regulation and the evolution of sophisticated chemical communication are fundamental to the transition to eusociality in insects. Strong changes in DNA methylation patterns correlated with broad-scale modifications of expression patterns. Many of these modified expression patterns remained consistent among the three studied termite species and occurred within protein pathways essential for eusocial life, such as CHC production, chemoperception, ecdysteroid synthesis and JH transport. The stronger patterns we observe for *M. natalensis*, especially within genes linked to chemical communication, such as the expansion of Desat B and CYP4G1 genes and significant positive selection in desaturases, elongases and in IRs, may be associated with this termite’s higher level of eusociality and its status as a superoganism.^13^ The analysis of further higher and lower termites would shed light on the generality of these patterns and possibly assist in the distinction between the influences of ecological and eusocial traits.

Many of the mechanisms implicated in the evolution of eusociality in the termites occurred convergently around 50 my later in the phylogenetically distant Hymenoptera. However, several details are unique due to the distinct conditions within which eusociality arose. One important difference is the higher TE content within cockroaches and termites, which likely facilitated changes in gene family sizes, supporting the transition to eusociality. However, the most striking difference is the apparent importance of IRs for chemical communication in the termites, compared to ORs in Hymenoptera. According to our results, the non-eusocial ancestors of termites possessed a broad repertoire of IRs, which favoured the evolution of important functions for colony communication in these chemoreceptors within the termites, whereas in the solitary ancestors of eusocial hymenopterans ORs were most abundant.^14, 25^ The parallel expansions of different chemoreceptor families in these two independent origins of eusociality indicate that convergent selection pressures existed during the evolution of colony communication in both lineages.

## METHODS

### Genome sequencing and assembly

Genomic DNA from a single *Blattella germanica* male from an inbred line (strain: American Cyanamid = Orlando Normal) was used to construct two paired-end (180 bp and 500 bp inserts) and one of the two mate pair libraries (2 kb inserts). An 8kb mate pair library was constructed from a single female. The libraries were sequenced on an Illumina HiSeq2000 sequencing platform. The 413 Gb of raw sequence data were assembled with Allpaths LG,^54^ then scaffolded and gap-filled using the in-house tools Atlas-Link v.1.0 (https://www.hgsc.bcm.edu/software/atlas-link) and Atlas gap-fill v.2.2. For *Cryptotermes secundus*, three paired-end libraries (250 bp, 500 bp and 800 bp inserts) and three mate pair libraries (2 kb, 5 kb and 10 kb inserts) were constructed from genomic DNA that was extracted from the head and thorax of 1 000 individuals, originating from a single, inbred field colony. The libraries were sequenced on an Illumina HiSeq2000 sequencing platform. The *C. secundus* genome was assembled using SOAPdenovo (v.2.04)^55^ with optimised parameters, followed by gapcloser (v1.10, released with SOAPdenovo) and kgf (v1.18, released with SOAPdenovo).

### Transcriptome sequencing and assembly

For annotation purposes, twenty-two whole body RNAseq samples from various developmental stages were obtained for *B. germanica*. For *C. secundus* RNAseq libraries were obtained for three workers, four queens and four kings, based on degutted, whole body extracts. In addition, we sequenced 10 *M. natalensis* RNAseq libraries from three queens, one king and six pools of workers. All libraries were constructed using the Illumina (TruSeq) RNA-Seq kit.

For protein coding gene annotation, *B. germanica* reads were assembled with *de novo* Trinity (version r2014-04-13).^56^ The *C. secundus* reads were assembled using i) Cufflinks on reads mapped with TopHat (version2.2.1),^57, 58^ ii) *de novo* Trinity;^56^ and iii) genome-guided Trinity on reads mapped with TopHat.

### Repeat annotation

A custom *C. secundus* and *B. germanica* repeat library was constructed using a combination of homology-based and *de novo* approaches, including RepeatModeler/RepeatClassifier (http://www.repeatmasker.org/RepeatModeler.html), LTRharvest/LTRdigest^59^ and TransposonPSI (http://transposonpsi.sourceforge.net/). The *ab initio* repeat library was complemented with the RepBase (update 29-08-2016)^60^ and SINE repeat databases, filtered for redundancy with CD-hit and classified with RepeatClassifier. RepeatMasker (version open-4.0.6, http://www.repeatmasker.org) was used to mask the *C. secundus* and *B. germanica* genome. Repeat content for the other studied species (Fig. 1) was obtained from the literature.^61, 62, 63, 64, 65, 66, 67^

### Protein-coding gene annotation

The *B. germanica* genome was annotated with Maker (version 2.31.8),^68^ using (i) the species-specific repeat library, (ii) *B. germanica* transcriptome data (22 whole body RNAseq samples), and (iii) the swissprot/uniprot database (last accessed: 21-01-2016) plus the *C. secundus* and *Zootermopsis nevadensis* protein sequences for evidence-based gene model predictions. AUGUSTUS (version 3.2),^69^ GeneMark-ES Suite (version 4.21)^70^ and SNAP^71^ were used for *ab initio* predictions. *Cryptotermes secundus* proteincoding genes were predicted using homology-based, *ab initio* and expression-based methods, and integrated into a final gene set (see Supplementary Material). Gene structures were predicted by GeneWise.^72^ The *ab initio* annotations were predicted with AUGUSTUS^73^ and SNAP,^71^ retained if supported by both methods and integrated with the homology-based predictions using GLEAN.^74^ Transcriptome-based gene models were merged with PASA^75^ and tested for coding potential with CPC^76^ and OrfPredictor.^77^ PASA gene models were merged with the homology-based and *ab initio* gene set, retaining the PASA models in case of overlap. Desaturases, elongases, chemosensory receptors, Cytochrome P450’s and genes involved in the juvenile hormone pathway were manually curated in Blattodea.

### Differential gene expression

The *C. secundus* and *M. natalensis* RNAseq libraries, were complemented with nine published *Z. nevadensis* libraries, yielding 2 to 6 libraries from workers, queens and kings for each termite. These were compared to six of the *B. germanica* libraries: two from 5*^th^* instar nymphs, two from 6*^th^* instar nymphs and two from adult females. Reads were mapped to the genome using HiSat2.^78^ Read counts per gene where obtained using htseq-count and DESeq2^79^ was used for differential expression analysis. Differential expression analysis between kings (M), queens (F) and workers (majors and minors combined for *M. natalensis*) was assessed for the termites. For *B. germanica* we evaluated the differential expression between adults and the two last nymphal stages combined, with the assumption that the final nymphal stages are homologous to termite workers and the adult females are homologous to termite queens. Genes were considered significantly differentially expressed if p < 0.05 and log_2_ fold change > *|*1*|* in order to account for allometric differences as recommended by Montgomery and Mank.^80^

### Protein orthology

In addition to *B. germanica*, *C. secundus*, *Z. nevadensis* and *M. natalensis*, 16 other insect proteomes were included in our analyses; *L. migratoria*, *R. prolixus*, *E. danica*, *D. melanogaster*, *A. aegypti*, *T. castaneum*, *N. vitripennis*, *P. canadensis*, *A. mellifera*, *H. saltator*, *L. humile*, *C. floridanus*, *P. barbatus*, *S. invicta*, *A. echinatior* and *A. cephalotes*; as well as for the centipede, *S. maritima*, as an outgroup (for sources see Supplementary Table 22). These proteomes were grouped into orthologous clusters with OrthoMCL,^81^ with a granularity of 1.5.

### IR and OR identification, phylogeny and structure

Ionotropic receptors (IRs) were identified using two custom Hidden Markov Models (HMMs) obtained with `hmmbuild` and `hmmpress` of the HMMER suite.^82^ The first HMM comprises the IR’s ion channel and ligand-binding domain based on a MAFFT^83^ protein alignment of 76 IRs from 15 species (Supplementary Table 23). The second HMM was built to distinguish IRs from iGluRs, IR8a and IR25a, which have an additional amino-terminal domain (ATD).^24^ For this we built an HMM from 48 protein sequences (Supplementary Table 23). The proteomes were scanned with pfam scan and the two custom HMMs, where proteins that matched the IR HMM, but not the ATD HMM were annotated as IRs. ORs were identified based on the Pfam domain `PF02949` (7tm Odorant receptor).

Multiple sequence alignments of IRs and ORs were obtained with hmmalign,^82^ using the Pfam OR `HMM PF02949` and custom IR HMM to guide the alignment. Gene trees were computed with FastTree^84^ (options: `‐pseudo ‐spr 4 ‐mlacc 2 ‐slownni`) and visualised with iTOL v3.^85^ Putative IR ligandbinding residues and structural regions were identified based on the alignments with *D. melanogaster* IRs and iGluRs of known structure.^86^

### Gene family expansions and contractions

For the analyses of gene family expansions and contractions, the hierarchical clustering algorithm MC-UPGMA^87^ was used, with a ProtoLevel cutoff of 80.^88^ Protein families were further divided into sub-families if they contained more than 100 proteins in a single species, or more than an average of 35 proteins per species. Proteins were blasted against the RepeatMasker TE database (E-value < 10*^−5^*) and clusters where > 50% of the proteins were identified as transposable elements were discarded. Clade- and species-specific protein family expansions and contractions, were identified with CAFE v3.0^89^ using the same protocol as^9, 10^ (see also Supplementary Material).

### TE-facilitated expansions

The repeat content in the 10 kb flanking regions of *B. germanica*, *C. secundus*, *Z. nevadensis* and M.*natalensis* genes was calculated using bedtools.^90^ CDS’ from neighbouring genes were removed and the repeat content was analysed using Generalized Linear Mixed Models (glmmPQL implemented in the R^91^ package MASS^92^) with binomial error distribution. Fixed predictors included gene family expansion, species ID and their interaction. Cluster ID was fitted as random factor to avoid pseudo-replication. Significance was assessed based on the Wald-*t* test (R package aod^93^) at *α* < 0.05. Main and interaction effects for each of the genomic regions are listed in table S8. Model parameters are listed in table S8.

### Tests for positive selection

To test for positive selection within gene families of interest, (i) site model tests 7 and 8 were performed (model = 0; NSsites = 7 8) on species-specific CDS alignments or ii) branch-site test (model = 2; NSsites = 2; fix omega = 1 for null model and 0 for alternative model) on multi-species alignments. Protein sequences were aligned using MAFFT^83^ with the E-INS-i strategy, and CDS alignments were created using pal2nal.pl.^94^ Phylogenetic trees were created with FastTree.^84^ Alignments were trimmed using Gblocks (settings: ‐b2 = 21; ‐b3 = 20; ‐b4 = 5; ‐b5 = a). Models were compared using LR test and where p < 0.05, Bayes Empirical Bayes (BEB) results were consulted for codon positions under positive selection.

### CpG depletion patterns and GO enrichment

To estimate DNA methylation we compared observed to expected CpG counts within CDS sequences.^38, 39^ A low CpG*_o/e_*indicates a high level of DNA methylation, as the cytosine of methylated CpGs often mutate to thymines. Expected CpG counts were calculated by dividing the product of cytosine and guanine counts by the sequence length. The PCA in figure 4 was created using the R function prcomp on log transformed CpG*_o/e_*values for all 1-to-1 orthologs for the seven hemimetabolous species. These orthologs were extracted from the OrthoMCL results. The 3D plot was created with the plot3d command from the R package rgl.

CpG depleted (first quartile) and enriched genes (fourth quartile) were tested for enrichment of Gene Ontology terms. Pfam protein domains were obtained for *B. germanica*, *Z. nevadensis*, *C. secundus* and *M. natalensis* protein sequences using PfamScan.^95^ Corresponding GO terms were obtained with Pfam2GO. GO-term over-representation was assessed using TopGO^96^ package in R. Enrichment analysis was performed using the weight algorithm selecting nodesize=10 to remove terms with less than 10 annotated GO terms. After that GO terms classified as significant (topGOFisher<0.05) were visualized using R package tagcloud (https://cran.r-project.org/web/packages/tagcloud/).

### Data availability

The data reported in this study are archived at the following databases: NCBI (genomes sequences), SRA (genomic and transcriptomic reads), i5k Workspace@NAL & Dryad (annotations). Detailed accession information is tabulated in the Supplementary Materials (Supplementary Table 24). Scripts and output files are available on request from E.B.B.

## Acknowledgements

We thank Oliver Niehuis for allowing use of the unpublished *E. danica* genome, Jürgen Gadau and Chris Smith for comments and advice on the manuscript, Jonathan Schmitz for assistance with analyses and proof-reading the manuscript. JK thanks Charles Darwin University (Australia), especially Prof. Stephen Garnett and the Horticulture and Aquaculture team for providing logistic support to collect *C. secundus*. The Parks and Wildlife Commission, Northern Territory, the Department of the Environment, Water, Heritage and the Arts gave permission to collect (Permit number 36401) and export (Permit WT2010-6997) the termites. USDA is an equal opportunity provider and employer. MCH and EJ supported by DFG grant BO2544/11-1 to EBB. JK by University of Osnabrück and DFG grant KO1895/16-1. XB and MDP supported by Spanish Ministerio de Econom´ıa y Competitividad (CGL2012-36251 and CGL2015-64727-P to XB, and CGL2016-76011-R to MDP), including FEDER funds, and by Catalan Government (2014 SGR 619). CS: grants from US Department of Housing and Urban Development (NCHHU-0017-13), National Science Foundation (IOS-1557864), Alfred P. Sloan Foundation (2013-5-35 MBE), National Institute of Environmental Health Sciences (P30ES025128) to Center for Human Health and the Environment, and Blanton J. Whitmire Endowment. MP is supported by a Villum Kann Rasmussen Young Investigator Fellowship (VKR10101).

## Author contributions

E.B-B. conceived, managed and coordinated the project; M.C.H., E.J. and H.M.R. are joint first authors. J.K. conceived and managed *C. secundus* sequencing project, coordinated termite-related analyses; S.R. conceived and managed *B. germanica* sequencing project; S.R., S.D., S.L.L., H.C., H.V.D., H.D., Y.H., J.Q., S.C.M., D.S.T.H., K.C.W., D.M.M. and R.A.G. carried out *B. germanica* library construction, genome sequencing and assembly; C.S., A.W.K. provided biological material through full-sib mating for B. *germanica*; X.B. and C.S. co-managed the *B. germanica* analysis; M.P. and C.P.C. implemented Web Apollo data traces; S.O. and M.P. provided biological material for *M. natalensis*; C.G., J.G., J.M.M.-K., A.M., F.S., H.H. & J.K. coordinated and carried out DNA and RNA sequencing for *C. secundus*; M-D.P., X.B. and G.Y. coordinated transcriptome sequencing of *B. germanica*; L.M. performed automated gene prediction on *C. secundus*; E.J. and N.A. improved assembly and annotation for *B. germanica* & C. *secundus*, compared and analysed genome sizes and quality. E.J., N.A. and L.P.M.K. analysed TEs; M.C.H. analysed CpG patterns and signatures of selection; T.B-F., E.J., C.K., L.P.M.K. and A.L-E. performed orthology and phylogenetic analyses; L.P.M.K., E.J., H.M.R. and M.C.H. analysed gene family evolution; A.L-E., E.J. and M.C.H. analysed transcriptomes and performed DE analyses; T.B.-F. and A.L-E. carried out orthoMCL clustering; H.M.R. corrected gene models for chemoreceptors; C.K. and E.J. for desaturases and elongases; A-K.H. and M.C.H. of Cytochrome p450s; E.B-B and M.C.H drafted and wrote the manuscript; X.B., M-D.P., J.K. contributed to sections of the manuscript; E.J., L.P.M.K., A.L-E., C.K., M.C.H. wrote and organized Supplementary Materials; L.P.M.K., N.A., A.L-E., M.C.H. and E.B-B. prepared figures for the manuscript. All authors read, corrected and commented on the manuscript.

